# Machine learning strategies to tackle data challenges in mass spectrometry-based proteomics

**DOI:** 10.1101/2024.05.02.592141

**Authors:** Ceder Dens, Charlotte Adams, Kris Laukens, Wout Bittremieux

**Affiliations:** Adrem Data Lab, Department of Computer Science, University of Antwerp, Middelheimlaan 1, 2020 Antwerpen, Belgium

**Keywords:** computational proteomics, deep learning, peptide property prediction

## Abstract

In computational proteomics, machine learning (ML) has emerged as a vital tool for enhancing data analysis. Despite significant advancements, the diversity of ML model architectures and the complexity of proteomics data present substantial challenges in the effective development and evaluation of these tools. Here, we highlight the necessity for high-quality, comprehensive datasets to train ML models and advocate for the standardization of data to support robust model development. We emphasize the instrumental role of key datasets like ProteomeTools and MassIVE-KB in advancing ML applications in proteomics and discuss the implications of dataset size on model performance, highlighting that larger datasets typically yield more accurate models. To address data scarcity, we explore algorithmic strategies such as self-supervised pretraining and multi-task learning. Ultimately, we hope that this discussion can serve as a call to action for the proteomics community to collaborate on data standardization and collection efforts, which are crucial for the sustainable advancement and refinement of ML methodologies in the field.

## Introduction

In the rapidly evolving field of bioinformatics, machine learning (ML) has become an indispensable tool for analyzing proteomics data [1]. Particularly for complex mass spectrometry (MS) data, the integration of ML into data processing workflows marks a significant advancement in our ability to interpret the data. Early adoption of ML techniques, such as the semi-supervised rescoring of peptide–spectrum matches (PSMs) introduced by Percolator [2], has significantly boosted the accuracy and sensitivity of spectrum annotation. These early implementations laid the groundwork for a broader integration of ML into proteomics, demonstrating its potential to enhance data analysis.

In recent years, the field has witnessed a surge in the application of the latest generation of neural network approaches, driving the incorporation of ML into virtually every element of the proteomics data analysis pipeline. As an example, a popular approach is to use predicted fragment ion intensities [3], [4], [5], [6], [7], liquid chromatography (LC) retention times (RTs) [8], [9], and ion mobility collisional cross section (CCS) values [10], [11], [12] to generate increasingly accurate simulated data and to use this information for PSM rescoring [12], [13], [14]. Another example is the development of several deep learning-driven *de novo* peptide sequencing tools that have recently emerged [15], [16], [17]. The rapid succession of these new approaches over the past years and the performance gains they achieve underscores the dynamic nature of the field and the potential of ML to transform proteomics data analysis.

Despite the proliferation of ML tools in proteomics, the diversity of ML algorithms, including various model architectures and training datasets, poses a significant challenge for critically evaluating and comparing the performance of these methods. The unique contributions of each tool are often obscured by confounding factors, making it difficult to discern their true benefits. This lack of transparency is further exacerbated by the fact that biological data, and by extension, MS data, are inherently messy and devoid of a ground truth. Consequently, further progress in computational MS may not stem solely from increasingly advanced ML architectures. There is a growing recognition that it is crucial to invest in high-quality, large-scale datasets for ML model training, as well as consistent evaluation data for objective ML model benchmarking [18], [19], [20].

The heterogeneity of MS applications necessitates a collaborative effort across the scientific community to steer data standards and foster data collection efforts. Whereas file format standardization efforts, spearheaded by the Human Proteome Organization’s Proteomics Standards Initiative [21], have been a tremendous benefit to the field, a similar approach for data standardization in bioinformatics and ML is so far lacking. We are convinced that such a collective approach is essential for overcoming the limitations posed by data scarcity and future proofing the development of robust ML models in the field.

To offer a starting point for this discussion, here, we will highlight commonly used datasets in the development of ML models for MS-based proteomics and describe the applications in which they have been utilized. Additionally, we will explore strategies for harnessing data for various ML tasks in proteomics and address challenges associated with data scarcity.

This survey will cover lessons learned from ML approaches that have proven successful, but also those that have not. The latter is particularly relevant, as the scientific literature often omits information on unsuccessful methodologies. Yet, understanding these failures is crucial for the community, preventing redundant efforts and fostering a more efficient path toward innovation. By focusing on the strategic use of data rather than solely on algorithmic advancements, we aim to chart a course for future research in computational MS that leverages the full potential of ML to extract valuable insights from proteomics data.

### Commonly used datasets for machine learning in proteomics

First, we will introduce several key datasets that are commonly used for peptide property prediction, including predicted fragment ion intensities, RTs, and CCS values. While ML can be applied to a broader range of proteomic challenges [22], the datasets highlighted in this section are pivotal for advancing our understanding and capabilities in specific areas such as MS/MS spectrum prediction and *de novo* peptide sequencing, which are some of the currently most studied ML applications in proteomics.

#### ProteomeTools

The ProteomeTools project consists of over one million peptides that have been synthesized and analyzed, including tryptic peptides covering all canonical human proteins [18], peptides containing 21 post-translational modifications (PTMs) [19], and non-tryptic peptides [23]. The synthesized peptides were combined into pools of about 1,000 peptides, spiked with 66 non-naturally occurring and 15 stable isotope labeled peptides for RT calibration. To avoid ambiguity, pools were designed in such a way that peptides do not have identical masses [18].

The tryptic dataset can be divided into three different subsets. First, it includes a ‘proteotypic’ set of 24,875 peptides covering 15,855 human Uniprot/SwissProt [24] annotated genes that are confidently and frequently identified with prior MS evidence available in ProteomicsDB [25], [26]. Second, it includes a ‘missing gene’ set of 140,458 peptides covering 4,818 genes which lacked confident experimental identification evidence in ProteomicsDB. Third, it includes a subset of the human SRMAtlas [27] containing 90,967 peptides mapping to 19,099 genes, covering both proteins with empirical evidence as well as ‘missing’ proteins [27]. Each peptide pool was subjected to LC using a Dionex 3000 HPLC system (Thermo Fisher Scientific) with C18 columns coupled online to an Orbitrap Fusion Lumos mass spectrometer (Thermo Fisher Scientific). Each peptide pool was first measured using a “survey” method. From this data RT constraints were generated for each pool and used for three subsequent LC-MS/MS measurements focussing on different acquisition types: “HCD” (higher-energy collisional dissociation), “IT” (ion trap), and “ETD” (electron-transfer dissociation) [18].

Additionally, about 5,000 synthetic peptides carrying 21 different modifications were generated by modifying four residues (lysine, arginine, proline, tyrosine). Tryptic peptides were selected based on previously successfully detected synthesized peptides containing these amino acids. The peptide length was limited to 7–20 amino acids and the modification site was never located at the C-terminus. Each peptide pool was subjected to LC using a Ultimate 3000 nano-HPLC coupled to an Orbitrap Fusion Lumos ETD mass spectrometer (Thermo Fisher Scientific). Each peptide pool was first measured using a “survey” run, followed by three subsequent LC-MS/MS measurements, comprising a total of 11 different fragmentation modes: “3xHCD”, “2xIT_2xHCD”, and “ETD” [19]. These peptides were also measured using an EASY-nLC 1200 (Thermo Fisher Scientific) system coupled to a hybrid TIMS-quadrupole TOF mass spectrometer (Bruker Daltonik timsTOF Pro) [11].

The non-tryptic peptides can be separated into four different sets according to their origin. The “HLA Class I” set, containing 168,688 peptides with 8–12 amino acids, was obtained from the Immune Epitope Database (IEDB) [28] and the study from Bassani-Sternberg et al. [29]. The “HLA Class II” set was taken from the SysteMHC Atlas project (release April 2017) [30] and contains 73,464 peptides with 10–25 amino acids. The “AspN” and “LysN” peptide sets, respectively containing 31,744 and 31,435 peptides with 7–25 amino acids, were derived from two studies employing the proteases for deep proteome studies [31], [32]. Each peptide pool was subjected to LC using a Dionex 3000 HPLC system (Thermo Fisher Scientific) with C18 columns coupled to an Orbitrap Fusion Lumos mass spectrometer (Thermo Fisher Scientific). Each peptide pool was first measured using a “survey” run, followed by three subsequent LC-MS/MS measurements comprising a total of 11 different fragmentation modes: “3xHCD”, “2xIT_2xHCD”, and “ETD”, covering different fragmentation settings (HCD, CID, ETD, EThcD, ETciD) and mass analyzers (FTMS: Orbitrap mass analyzer, ITMS: ion trap mass analyzer) [23]. The same peptides were also measured using an Evosep One HPLC system (Evosep) coupled to a hybrid TIMS-quadrupole TOF mass spectrometer (Bruker Daltonik timsTOF Pro) [33].

ProteomeTools also includes a TMT subset, containing both tryptic and non-tryptic synthesized peptides labeled with a TMT 6-plex label. Peptide pools were subjected to LC using a Dionex 3000 HPLC system (Thermo Fisher Scientific) with C18 columns coupled to an Orbitrap Fusion Lumos mass spectrometer (Thermo Fisher Scientific), after which each pool was measured twice, covering seven different fragmentation modes: “TMT1” and “TMT2” [34].

To assign ground truth peptide labels to the MS/MS spectra, raw data were analyzed using MaxQuant [35], searching individual LC-MS/MS runs against pool-specific databases. In addition to filtering PSMs at a 0.01 posterior error probability (PEP), subset-specific Andromeda score cutoffs were applied to safeguard correct identification.

The ProteomeTools data consists of ground truth fragment ion intensities, RTs, and in some cases CCS values, that are expected to be largely free of interference due to the construction of the peptide pools. These data have been used to develop several ML tools, including the Prosit deep neural network for fragment ion intensity and RT prediction [4], and several *de novo* peptide sequencing tools [36], [37], [38], [39]. Additionally, ProteomeTools is often used as a benchmark dataset to evaluate prediction tools, such as the CCS prediction model from Meier et al. [11].

An important consideration to take into account is that while the interference-free construction of the peptide pools in ProteomeTools is instrumental in assigning ground truth peptide labels, at the same time it also results in MS/MS spectra that are more “clean” than those obtained from complex mixtures. As such, care has to be taken during ML development that the model can generalize to noisier MS/MS data as well. Furthermore, as all cysteines in ProteomeTools are carbamidomethylated, the data cannot be used to develop ML models that can be applied to peptides containing free cysteine side chains.

The ProteomeTools data are made publicly available through various datasets in the PRIDE repository [40] (Supplementary Table 1). Additionally, a subset of the data has been released as data frames, explicitly geared towards ML researchers [41].

#### MassIVE-KB

The MassIVE Knowledge Base (MassIVE-KB) [42] was compiled via secondary analysis of 31 terabytes of human HCD data, derived from over 669 million MS/MS spectra contained in 227 publicly accessible datasets. Peptide identification was performed using a consistent bioinformatics pipeline including sequence database searching with MS-GF+ [43] against the UniProt human reference proteome. Variable modifications that were considered include oxidation on methionine, N-terminal acetylation, carbamylation, pyro-glu on glutamine, and deamidation on asparagine and glutamine. Cysteine carbamidomethylation was specified as a fixed modification. A dynamic search space adjustment was performed for synthetic samples coming from the tryptic subset of the ProteomeTools project (see above) and the BioPlex affinity purification MS experiments [44], [45], reducing the protein database to only the relevant candidates to improve the sensitivity of those searches.

A key aspect during repository-scale spectrum annotation is that the false discovery rate (FDR) has to be carefully controlled. MassIVE-KB used a 1% PSM-level FDR for the initial spectrum annotations, resulting in 191,152,777 confidently annotated PSMs. Next, high-quality MS/MS spectra were selected based on the top 100 PSMs for each peptidoform (i.e. the unique combination of a peptide with its PTMs), with a limit of 20 spectra per original dataset to avoid bias. The peptidoforms were grouped by peptide length and filtered to a 1% local FDR using a sliding window of 500 peptidoforms, resulting in a global peptidoform-level FDR of 0.1%. Finally, the picked protein FDR strategy [46] was used to filter the results to 1% protein-level FDR.

Considering the remaining top 100 PSMs per peptidoform resulted in a subset of 30,160,134 PSMs with uniformly 0% q value in each PSM’s original search. To select representative MS/MS spectra, within each peptidoform all pairwise cosine similarities were calculated and the medoid spectrum, i.e. the spectrum with the highest average similarity to all other spectra, was selected. This process resulted in a spectral library containing 2,154,269 unique peptidoforms and 1,114,503 unique peptides, mapping to 19,611 human proteins. The final spectral library can be downloaded from the MassIVE-KB Peptide Spectral Libraries web page^1^ in the MGF and sptxt formats.

Given its high-quality MS/MS spectra and large size, MassIVE-KB has been used to train several ML applications, either using the final spectral library of 2.1 million MS/MS spectra, or the 30 million PSMs prior to spectrum deduplication. One of the first ML applications using MassIVE-KB was GLEAMS [47], which uses deep contrastive learning to convert MS/MS spectra to vector embeddings for subsequent clustering. Additionally, MassIVE-KB has been used to train several *de novo* peptide sequencing [17], [38], [48] and fragment ion intensity prediction models [49].

There are a few considerations to take into account when using MassIVE-KB that mirror those mentioned previously. First, MassIVE-KB used a multi-tiered process for stringent FDR control. While this is essential to avoid propagation of incorrect spectrum assignments and ensures that the spectra are of high quality, similar to the ProteomeTools data, this might not be entirely representative of typical MS/MS data that could exhibit higher noise levels. Additionally, MassIVE-KB only contains peptides with carbamidomethylated cysteines, considered only a limited set of the most common chemically introduced PTMs, and contains HCD data from Orbitrap instruments only.

A newer version of the MassIVE-KB spectral library has since been released and is available from the corresponding web page. This MassIVE-KB v2 spectral library contains 5,948,126 MS/MS spectra from 2,492,495 unique peptides, covers a wider range of PTMs, as well as multi-enzyme data. However, as it has currently not been described in a scientific paper yet (April 2024), the precise data origins and bioinformatics workflow to obtain these results has not been detailed yet.

#### Chronologer

Chronologer [50] is a residual convolutional neural network developed for peptide RT prediction. To train their prediction tool, the authors undertook significant efforts in composing a large-scale, comprehensive peptide RT dataset by aggregating and harmonizing 11 public datasets. Given that each of these peptide datasets were collected using a different HPLC system, C18 resin, and LC gradient, individual RTs between datasets are not directly comparable and needed to be aligned into a common RT space for comparison. Due to the necessity of sufficient dataset overlap for nonlinear regression, the RT alignment was performed using predicted unitless relative RTs from Prosit [4], using a kernel density estimation-based alignment algorithm [51]. This approach demonstrated robustness to outliers from incorrect peptide identifications as well as correlated errors in Prosit predictions [52].

The final dataset consists of over 2.6 million peptides and is available on GitHub^2^. It includes both tryptic and non-tryptic peptides, containing 13 different PTMs, as well as unmodified peptides. The data harmonization efforts conducted are highly relevant, resulting in a final dataset that exceeds any individual study. Nevertheless, during our analysis, we identified some inconsistencies and potential quality issues, indicating the inherent complexities of combining data from heterogeneous sources. After addressing these issues by removing certain peptides, the final dataset comprised 2.1 million unique peptides containing 12 PTMs (see Methods section for details).

While the datasets discussed are regularly utilized in the proteomics field for ML applications, it is important to recognize that there are other valuable datasets which also play a crucial role in the development and testing of ML methods. Notable among these are the NIST peptide spectral libraries [53], which were one of the earliest sources of “ground truth” MS/MS data. Additionally, a nine-species benchmarking dataset [15], originally compiled by the DeepNovo authors, is commonly used to train and evaluate *de novo* peptide sequencing tools. Finally, resources like MassIVE.quant [54] and quantms [55] can help support the emerging development of ML solutions for peptide and protein quantification. These datasets, though not covered in detail in this section, are highly recommended for researchers looking to explore new avenues in ML within proteomics, underscoring the dynamic and ever-expanding nature of this field.

### Machine learning performance is driven by data availability

Although ML has spurred significant progress in MS, current models often solely rely on the information provided in their training data, lacking broader contextual understanding or expert knowledge. Consequently, there is an increasing need for expansive and comprehensive datasets that capture the full spectrum of potential inputs. This requirement is especially critical for deep learning models, given their significantly higher complexity.

To gain insight into the current state of peptide property prediction datasets, we evaluated the performance of a transformer neural network encoder [56] in predicting the RT and CCS of a mixture of modified and unmodified peptides that were not previously seen. For RT prediction, we used a curated version of the Chronologer dataset [50], containing 2.1 million peptides and 12 PTMs. The dataset compiled by Meier et al. [11], featuring ∼714,000 peptides and 3 PTMs, was used for CCS prediction. The models were trained on various subsets of the data. For both RT and CCS predictions, we initially reserved 10% of the data for testing and another 10% validation. The remaining data was randomly subsampled to create training datasets comprising 5%, 10%, 25%, 50%, 75%, and 100% of the training data.

The transformer architecture, renowned for its excellent performance not only on text-related tasks but also on various protein-related tasks, served as the backbone of our models. Previous research by Pham et al. [57] showed the effectiveness of the transformer architecture in RT prediction. In our experiments, we fine-tuned the TAPE [58] transformer, which is pretrained using self-supervised learning on a large-scale protein dataset.

Our analysis reveals a decrease in both RT and CCS prediction errors as the dataset size increases (Figure 1). Measuring performance with the Δ95% or Pearson correlation coëfficient gives similar results (Supplementary Figure S1). The RT prediction error shows a clear downward trend, indicating that further expansion of the training data could significantly enhance the accuracy of the model’s predictions. In contrast, while the CCS prediction error appears to level off after reaching 75% of the current dataset size, seemingly indicating that maximal performance has been achieved, we believe that in this case additional data could still yield further improvements. First, it is important to acknowledge that only one test was conducted per training data fraction. As a result, the performance of the 50%, 75%, and 100% data fractions might deviate from their average performance. Therefore, despite the current trend indicating a plateau in performance, it is plausible that further augmentation of the dataset could still lead to enhanced prediction results. Additionally, it is important to note that the CCS dataset encompasses only three distinct PTMs, in contrast to the iRT dataset, which includes 12 different PTMs and even more amino acid–PTM combinations. To broaden the applicability of CCS prediction models, it is crucial for them to perform accurately across a wide range of PTMs. As this would substantially expand the potential input space, we hypothesize that maintaining good performance will necessitate a proportional increase in the training dataset size.

**Figure 1.**
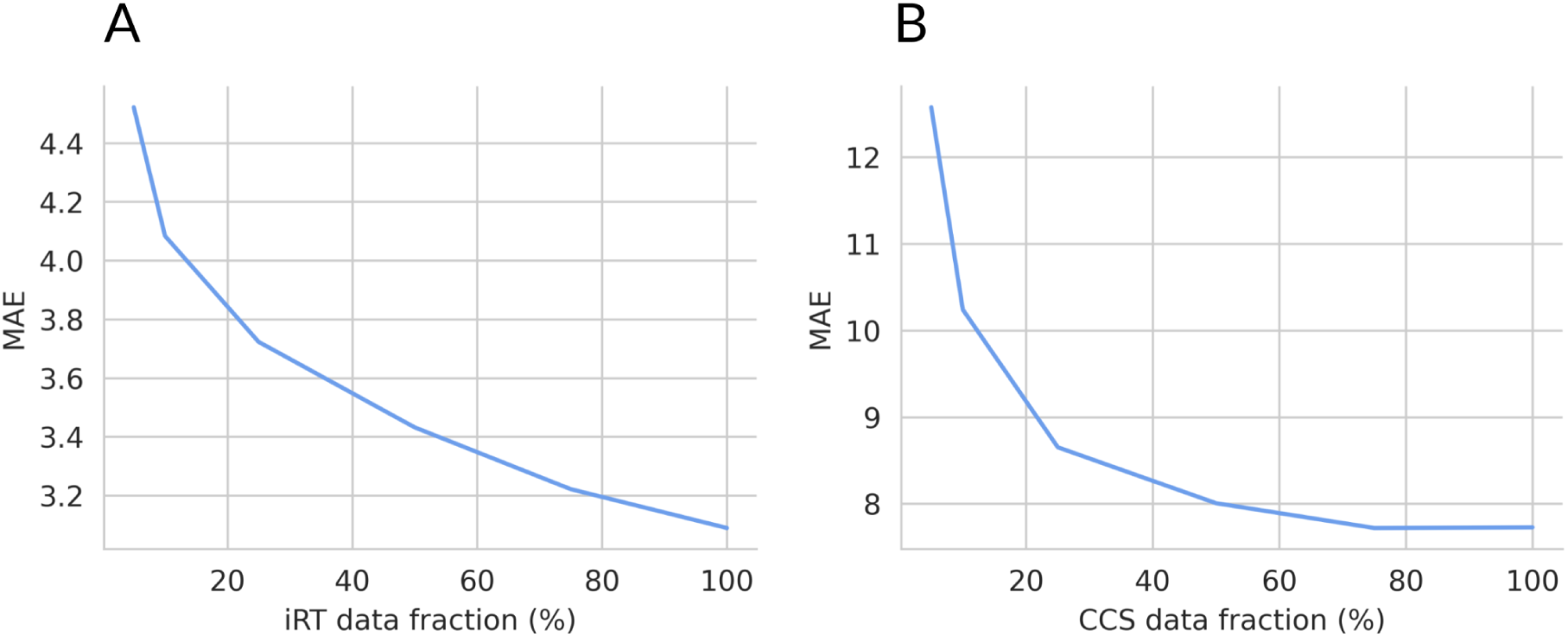
Peptide property prediction learning curves. The performance of a transformer encoder when trained on increasing data sizes. The performance is measured by the mean absolute error (MAE) of the model predicting the **(A)** indexed RT (iRT) and **(B)** CCS of unseen, optionally modified peptides. 100% of the RT data corresponds to around 1.7 million peptides and 100% of the CCS data is around 539,000 peptides.

### Machine learning strategies to mitigate data scarcity

In addition to algorithmic improvements and expanding dataset sizes, various techniques can reduce the impact of data scarcity in ML. Many of these approaches involve leveraging additional data, whether unlabeled or from a related task. Common examples include self-supervised pretraining followed by supervised fine-tuning, transfer learning, and multi-task learning.

We investigated the effects of self-supervised pretraining, using both additional related data and the same dataset, as well as multi-task learning on peptide property prediction performance. For the following experiments, we used a combination of the RT dataset, which is a curated version of the dataset composed by Chronologer [50], and the dataset composed by Meier et al. [11] for CCS values.

#### Pretraining improves model performance

When labeled data of high quality is scarce, self-supervised pretraining is a valuable and widely applied approach to enhance the performance of ML models. This method can utilize the same data employed for the model’s supervised learning task or a significantly larger unlabeled dataset. The objective of these methods is for the model to better grasp the structure and features of the input data, potentially resulting in accelerated convergence and improved performance upon fine-tuning with the actual task data.

Using a different dataset often offers the advantage of having access to a more extensive pool of data, aiding the model in gaining a comprehensive understanding of the entire potential input space and enabling it to learn more complex patterns. However, a potential caveat is that when the additional data substantially differs from the task data, pretraining might have minimal effects.

Self-supervised pretraining of transformer encoder models typically involves replacing the last few layers of the model, which predict label values, with a masked language model head [56]. In this approach, a small fraction of the input tokens are masked, and the model is trained to predict the tokens originally on these positions. This process does not require any labeled data, making it a self-supervised pretraining technique.

To assess the impact of pretraining, we conducted a comparative analysis using our multi-task transformer model trained with three distinct strategies on a five-fold cross-validation dataset comprising peptides with RT and CCS values. The first strategy involved no pretraining. In this approach, the model weights were randomly initialized and optimized solely using labeled task data. The second strategy consisted of pretraining followed by fine-tuning on the same dataset. We performed self-supervised pretraining with a masked language head using the training datasets from the cross-validation splits without labels. Subsequently, supervised fine-tuning of the model with task heads was conducted using the same dataset, learning the task labels. The third consisted of fine-tuning a model pretrained on a large protein dataset. We initialized the model with the weights of TAPE [58], pretrained on 31 million protein domains from Pfam [59], with only the weights of the multi-task prediction heads initialized at random. Using the pretrained TAPE transformer has already demonstrated beneficial performance for fragment ion intensity prediction [60]. Here, we used the pretrained TAPE model for fine-tuning using the cross-validation training datasets. By comparing the performance of the model under these three strategies, we aimed to elucidate the efficacy of different pretraining approaches in enhancing peptide property prediction accuracy.

Although the goal of this study is not to introduce the transformer encoder as the state-of-the-art peptide property prediction architecture, it is important to acknowledge that the reference models, Prosit for RT prediction and DCCS for CCS prediction, outperform the transformer when no pretraining is employed (Figure 2).

**Figure 2.**
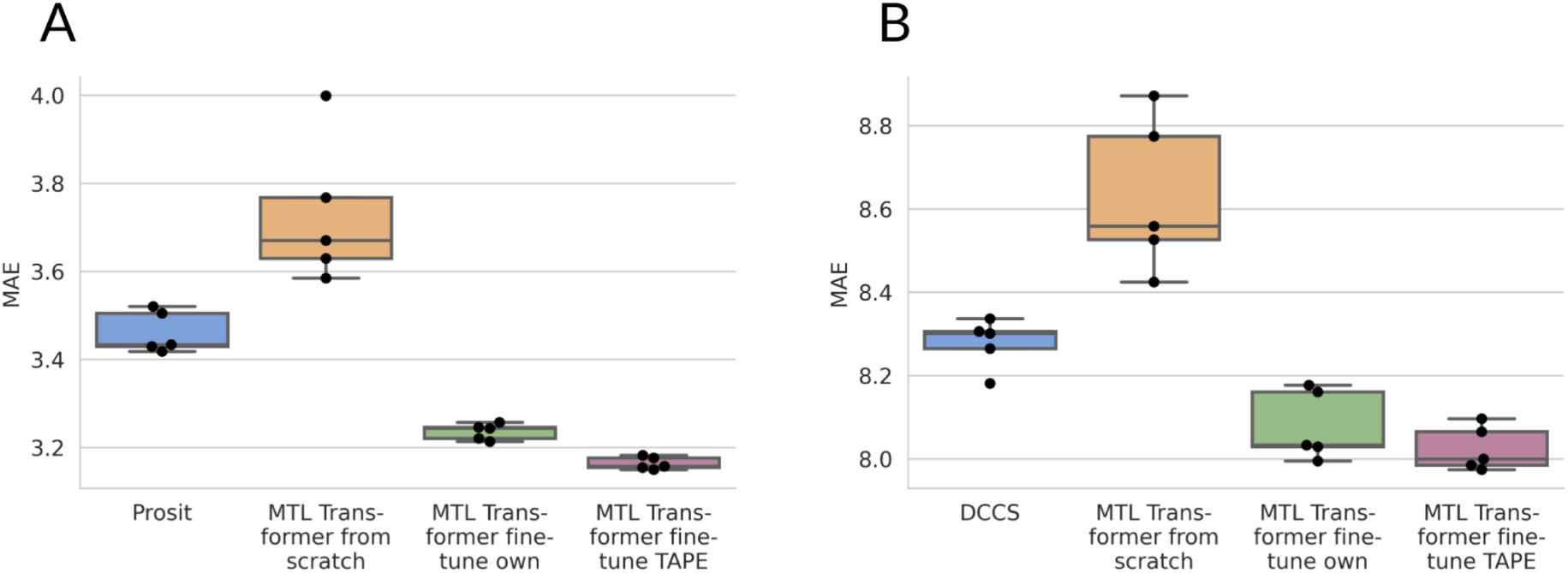
Influence of pretraining on peptide property prediction performance. The five-fold cross-validation performance of multi-task learning (MTL) transformer encoders when trained without pretraining (from scratch), when fine-tuning a model pretrained on the same data (fine-tune own), or when fine-tuning a model pretrained on a large protein dataset (fine-tune TAPE). The performance of a single cross-validation split is measured by the mean absolute error (MAE) of the model predicting the **(A)** iRT and **(B)** CCS of unseen peptides. The performance of **(A)** Prosit [4] for iRT prediction and **(B)** DCCS [11] for CCS prediction, retrained and evaluated on the same cross-validation splits, is also given as reference. The difference in performance of all models was tested with the Mann-Whitney U test, with the null hypothesis that the MAEs of the five-fold cross validation are from the same distribution. Making pairwise comparisons of all models, we found that their performance is all significantly different (Mann-Whitney U test, p = 0.008, two-tailed), except for the CCS prediction performance of the fine-tune own and fine-tune TAPE models (Mann-Whitney U test, p = 0.310, two-tailed). Boxplots are constructed as follows: the box extends from the lower to upper quartile values of the data, with a line at the median. The whiskers extend from the box to the last datum before 1.5 times the interquartile range above/below the box.

Our primary interest was exploring the potential of pretraining deep learning models to improve peptide property prediction performance. Our analysis illustrates that applying any type of pretraining proves beneficial for the transformer model, resulting in reductions in both mean absolute error and its variation, thus achieving more accurate and consistent models. For peptide RT prediction, fine-tuning a model pretrained on a large protein dataset, i.e. TAPE, yields even statistically significantly better results, while this is not the case for CCS prediction models. The results when measuring performance with the Δ95% and Pearson correlation coëfficient can be found in Supplementary Figure S2.

These results underscore the substantial improvement in performance achievable through self-supervised pretraining of deep learning models, even without additional data. They highlight the models’ capacity to discern certain features or patterns within the input data that might be overlooked when relying solely on supervised learning.

The enhanced performance of the RT prediction model when using a large, general protein dataset shows the effectiveness of self-supervised learning with slightly dissimilar data. This suggests that the high volume and diversity of the self-supervised data are more important than its similarity to the task-specific data. For CCS prediction, we do not see a significant difference between pretraining with the same data and fine-tuning the pretrained TAPE weights. This suggests that the lower variability in the peptide sequences of the CCS dataset cannot benefit from the high variability of the external pretraining dataset.

#### Multi-task learning fails to improve model performance

Another method to overcome the effects of limited ground truth data is multi-task learning [61]. This approach aims to leverage labeled data from a different task that shares similar input data. Like pretraining, multi-task learning can benefit from exploring a larger portion of the input space, finding previously unclear patterns. Moreover, multi-task learning has the potential to reduce dataset specific biases. By combining data from multiple datasets with varying noise and biases, the model becomes more adept at discerning the underlying signal. Additionally, when tasks are sufficiently similar, the model can benefit from signals that determine other task labels for its own task.

When using multi-task learning with deep learning models, we can distinguish two different methods: hard and soft parameter sharing [61]. With hard parameter sharing, a model is constructed in such a way that a portion of its layers are shared among multiple tasks. This part of the model creates a shared embedding of the input data, which is subsequently used to predict task-specific labels via designated task heads. Conversely, soft parameter sharing duplicates the model for each task, while linking a portion of the model’s weights to inhibit them from becoming too dissimilar. This allows the embeddings for different tasks to diverge, while still facilitating information transfer among them.

To investigate the effects of multi-task learning on peptide property prediction models, we performed a five-fold cross-validation comparison between a multi-task and a single-task transformer encoder for predicting peptide RT and CCS. For the single-task model, we use the TAPE model, pretrained on a large protein dataset, which was then fine-tuned separately using peptide RT and CCS data. The multi-task learning model is also based on the TAPE transformer and was pretrained on the same large protein dataset. However, in this model, the head responsible for predicting a single task label was replaced with multiple heads, each dedicated to a specific task (Figure 3), making this a hard parameter sharing multi-task learning model. The multi-task learning model was fine-tuned using a combination of labeled peptide RT and CCS data.

**Figure 3.**
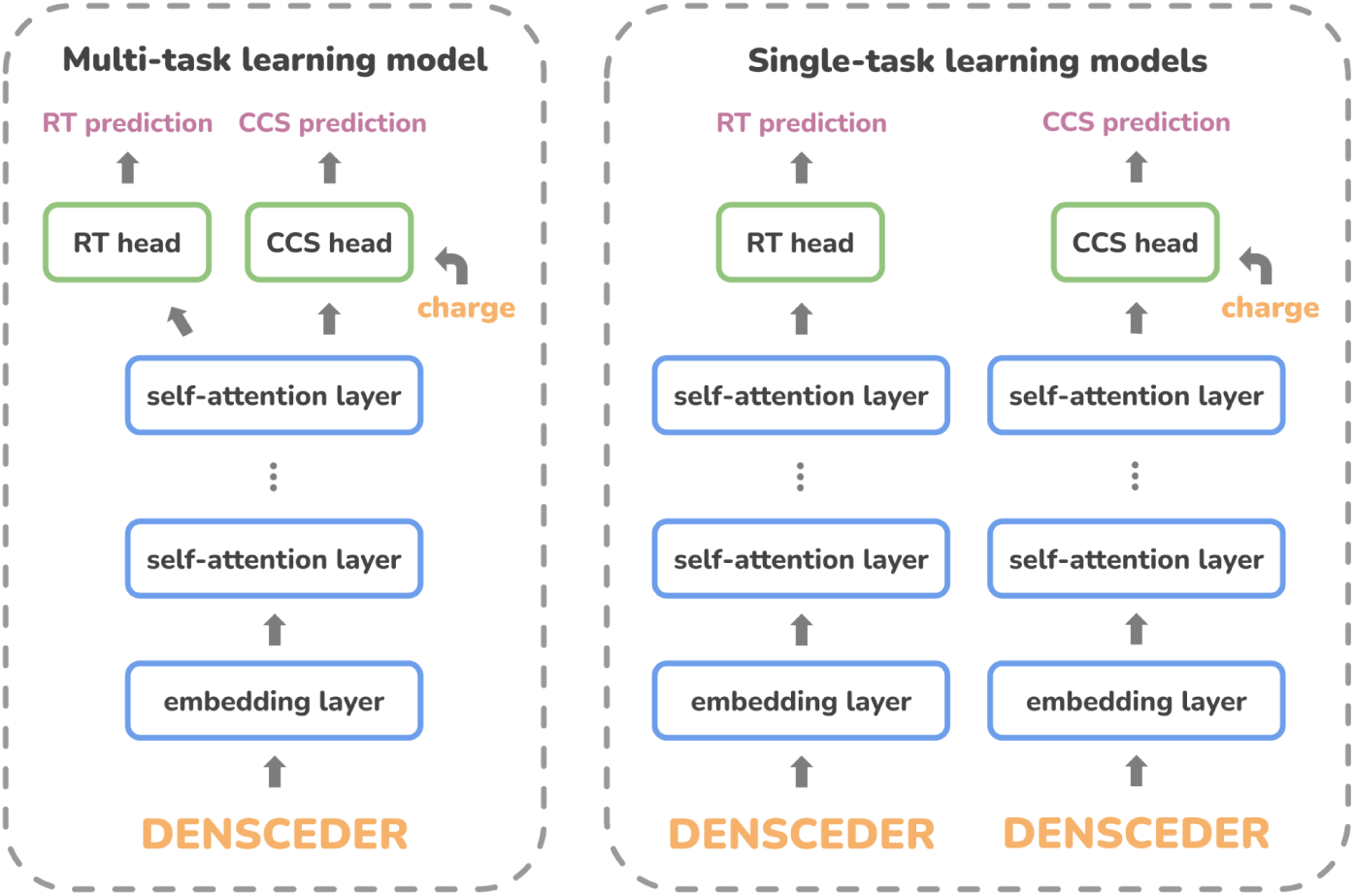
Schematic overview of the peptide property prediction models. A high-level overview of the components involved in the multi-task (left) and single-task (right) learning models. The inputs consist of the peptide sequence, which may contain PTMs, along with the precursor charge. Deep learning layers are shown as rounded rectangles and at the end, RT and CCS values are predicted.

Our experiments indicate that multi-task learning does not enhance the performance of peptide property prediction models compared to single-task learning (Figure 4 and Supplementary Figure S4). When comparing RT prediction error (Figure 4a), both models achieve a similar performance, with neither significantly outperforming the other (Mann-Whitney U test, p = 0.310, two-tailed). One possible explanation is the substantial size difference between the training datasets for RT prediction and CCS prediction. Consequently, the addition of CCS data might have a relatively minor impact. Moreover, the CCS dataset only contains three PTMs, which are already well-represented in the RT dataset. As a result, the CCS data may not contribute significantly to the RT prediction task, thus failing to enhance performance over the single-task learning model.

**Figure 4.**
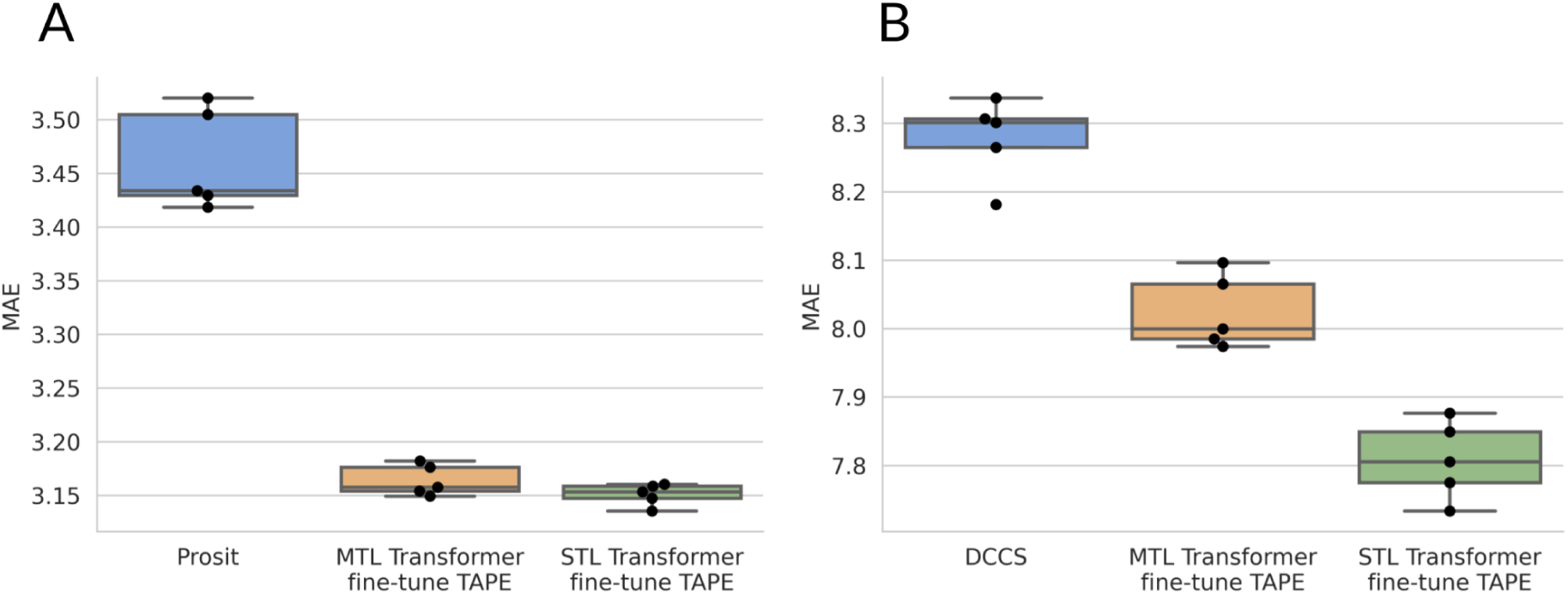
Influence of multi-task learning on peptide property prediction performance. The five-fold cross-validation performance of a multi-task learning (MTL) and a single-task learning (STL) transformer encoder. The performance of a single cross-validation split is measured by the mean absolute error (MAE) of the model predicting the **(A)** iRT and **(B)** CCS of unseen modified peptides. The performance of **(A)** Prosit [4] for iRT prediction and **(B)** DCCS [11] for CCS prediction, retrained and evaluated on the same cross-validation splits, is also given as reference. Boxplots are constructed as follows: the box extends from the lower to upper quartile values of the data, with a line at the median. The whiskers extend from the box to the last datum before 1.5 times the interquartile range above/below the box.

Interestingly, we also observe that the CCS prediction multi-task learning model gets significantly higher errors than the single-task model (Figure 4b) (Mann-Whitney U test, p = 0.004, one-sided). This indicates that adding RT data worsens CCS prediction performance. Several factors could explain this phenomenon. First, the CCS dataset is substantially smaller than the RT dataset, potentially leading to overfitting of the multi-task model on RT data, thereby diminishing performance on CCS data. Second, both the training and test datasets for CCS prediction have a low PTM variability. The additional variability introduced by the RT data might confuse the multi-task learning model, introducing irrelevant information for the current CCS test dataset. We hypothesize that additional CCS data with more PTM variability could rectify these disparities and would unlock the potential of multi-task learning.

## Conclusion

Recent years have seen a proliferation of ML applications, especially with the arrival of neural network approaches. These advancements span various facets of proteomics analysis, from predicting fragment ion intensities and RTs to developing deep learning-driven *de novo* peptide sequencing tools. Such rapid progress underscores ML’s potential to reshape proteomics data analysis profoundly.

Despite these technological strides, challenges remain in comparing the performance of ML tools due to the diversity of model architectures and training datasets. The absence of standardized data further complicates the evaluation process, emphasizing the need for high-quality, large-scale datasets that are essential for effective benchmarking. While there have been significant strides in the field to standardize file formats, similar efforts in data standardization for computational MS are still lacking. Collective action within the scientific community is crucial to address these limitations and to foster the development of robust ML models.

Our study underscores the importance of expansive datasets that include a diverse array of peptides and PTMs to train ML models effectively. We have demonstrated how increasing the dataset size enhances model performance, thereby reinforcing the need for comprehensive data resources. In an effort to mitigate the impact of data scarcity, we explored various techniques, such as self-supervised pretraining and multi-task learning, for peptide property prediction. Our findings reveal that while masked language model pretraining on peptide sequences can significantly improve model performance, multi-task learning did not provide substantial benefits. Although conceptually promising, multi-task learning requires highly similar tasks and sufficient data variability and scale across all tasks to be effective.

Transfer learning is another related approach that we briefly touched upon, which has shown promise in various domains [58], [62], [63], [64]. These methods essentially combine pretraining and multi-task learning, enabling the transfer of knowledge from one task to another. As an example, transfer learning has enabled the adaptation of models like Prosit to predict fragment ion intensities for different types of MS data [33].

Finally, we would like to call upon the proteomics community to prioritize the generation, compilation, and curation of datasets. While the development of new ML models is often more highlighted, the community stands to benefit more substantially from robust dataset creation and rigorous benchmarking. We encourage researchers to focus on these foundational efforts before pursuing the development of complex, multifaceted models, which often lack a standardized framework for evaluation. This strategic focus will likely accelerate scientific discovery and facilitate the development of more accurate and generalizable ML tools in proteomics.

## Methods

### Data

The experimental data used in this study comprised two distinct datasets: one containing RT measurements and the other containing CCS values. The multi-task learning models were trained and evaluated using the joint datasets. Duplicate modified peptide sequences were only removed per task, as further explained in subsequent sections, but not overall. When partitioning the data in training, validation, and testing sets, peptides with the same unmodified amino acid sequence were always assigned to the same split.

#### Peptide retention time data

The Chronologer dataset [50] was used as the reference dataset for the RT prediction tasks. This dataset consists of more than 2.6 million peptides from various sources, containing tryptic peptides, immunopeptides, as well as peptides originating from multiple organisms. This dataset features peptides with 13 different PTMs, present on various amino acids. Due to potential variations from different experimental conditions, such as different instruments, different labs, and experiments performed on different days, the RT has to be standardized to iRT. This calibration process was already applied for the Chronologer dataset.

To further ensure data integrity, quality, and consistency, several data curation steps were undertaken. First, all peptides with an iRT of more than 150 were deemed anomalous as they fall outside of the chromatographic range [65] and were removed, eliminating around 132,000 peptides. Subsequently, we found recurring iRT values across numerous distinct peptides. These stem from technical or post-processing artifacts and we removed all peptides associated with iRT values appearing 300 times or more from the same source, resulting in the removal of around 35,000 peptides. Additionally, peptides featuring PTMs deemed impossible, such as M acetylation and Q acetylation, were discarded, amounting to approximately 1,400 peptides. Lastly, duplicate peptides were merged by computing their median iRT values, resulting in the loss of around 367,000 peptides. This curation process resulted in an RT dataset comprising 2.1 million unique peptides containing 12 PTMs.

#### Peptide collisional cross section data

The dataset collected and analyzed by Meier et al. [11] was used for the CCS prediction tasks. We merged their training dataset, containing around 560,000 peptides, with their external test dataset, containing around 154,000 peptides. This resulted in a final dataset containing around 714,000 unique peptide–charge pairs, featuring three different PTMs (cysteine carbamidomethylation, methionine oxidation, and N-terminal acetylation). These peptides originated from whole-cell proteomes of *Caenorhabditis elegans*, *Drosophila melanogaster*, *Escherichia coli*, *HeLa*, and *Saccharomyces cerevisiae*, using up to three different enzymes with complementary cleavage specificity (trypsin, LysC, and LysN) [11]. Given that the precursor charge significantly influences CCS measurements, duplicate peptides with distinct charge states were retained in the dataset. Specifically, 67.8% of the peptides featured a charge state 2, while 27.0% exhibited a charge state of 3, and 5.2% possessed a charge state of 4.

#### Learning curve data

To construct learning curves, models were trained on different subsets of the RT and CCS datasets and afterwards evaluated. Initially, a random split was performed to generate testing and validation dataset splits, which remained consistent throughout subsequent analyses. After adding peptides sharing identical unmodified amino acid sequences, both test and validation datasets comprised 10% of the peptides from the complete dataset. Subsequently, from the remaining 80% of the data, 6 different RT and 6 different CCS training datasets were created by sampling 5%, 10%, 25%, 50%, 75%, and 100% of the RT and the CCS datasets.

#### Cross-validation data splits

The influence of pretraining and multi-task learning on model performance was assessed through five-fold cross-validation. The data splits were generated once and utilized for all different models. For the single-task models, only peptides with a label for the corresponding task were retained. First, the RT and CCS datasets were merged without removing or merging duplicates. Subsequently, a part of this dataset was randomly selected and all peptides sharing identical unmodified amino acid sequences were also added, resulting in a test dataset containing approximately 20% of the peptides of the full dataset. From the remaining 80%, another part was randomly sampled and peptides sharing identical unmodified amino acid sequences were added again, resulting in a validation dataset of around 10% of the full dataset. The remaining peptides, around 70% of the complete dataset, were used as the training dataset. This process was repeated five times, resulting in five distinct random training, validation, and testing cross-validation splits.

### Models

#### Transformer encoder

The single-task and multi-task models used in this study adopt a transformer encoder-only neural network architecture [56]. For our application, each amino acid–PTM combination is represented as a unique token, which are encoded into embeddings along with positional information using a trainable embedder. The input embedding for the first transformer layer is obtained by summing the token embedding and positional embedding. Subsequently, these embeddings go through a sequence of self-attention transformer encoder layers. The output of the final layer is pooled by extracting the embedding of the first token, followed by a fully connected layer.

The model’s architecture is different depending on whether it is undergoing pretraining or fine-tuning. During pretraining, a masked language model head is used. For this, a portion of the tokens in the input sequence are masked, and the masked language model head predicts the tokens at these masked positions using two fully connected layers, giving the probability for each token at each position.

For supervised learning of one or multiple task labels, each task gets its own task head. These heads receive the pooled output of the final self-attention transformer encoder layer and consist of a two-layer perceptron, predicting either the peptide iRT or CCS value. Notably, the CCS prediction heads also receive the one-hot encoded precursor charge as an additional input (Figure 3).

Furthermore, the transformer encoder for peptide property prediction standardizes the training data labels per task, ensuring that the labels get values with a mean of 0 and a standard deviation of 1. A reverse transformation is applied before outputting the validation and test predictions. This normalization procedure enhances model convergence and performance stability across tasks with different value ranges.

#### Prosit

Prosit [4] was used to compare RT prediction performance. The default Prosit model was trained from scratch using the same cross-validation data splits as the transformer encoder models.

#### DeepCollisionalCrossSection

To compare CCS prediction performance, a deep learning model using bidirectional LSTM units [11] was trained and tested using the same cross-validation data splits. Consistent with the name used in their implementation, we refer to this model as DeepCollisionalCrossSection (DCCS). Training of this model was done from scratch with the default model parameters.

#### Hyperparameter tuning of the transformer encoder

Transformer encoders have numerous tunable hyperparameters. However, certain parameters were fixed across our experiments. Specifically, we maintained a batch size of 1024 and a sequence length of 50. Four distinct model types were considered: training from scratch, pretraining, fine-tuning of self-pretrained weights, and fine-tuning of weights previously pretrained by TAPE. Each model type had multiple hyperparameters to tune, so SMAC3 [66] for fast hyperparameter optimization was used to make this feasible.

For the models trained from scratch, the hyperparameters that were optimized are the learning rate, the optimizer selection, the scheduler configuration, the embedding size, and the number of self-attention layers. In contrast, for the models undergoing pretraining or fine-tuning, we only optimized the learning rate, the optimizer selection, and the scheduler configuration. For the models undergoing pretraining, the embedding size and the number of self-attention layers were inherited from the best-performing model trained from scratch. Similarly, for models undergoing fine-tuning, these settings were inherited from the pretrained model.

The ranges considered for the hyperparameters were as follows: the learning rate spanned [1e-6, 1e-1] on a logarithmic scale; the embedding size ranged from 24 to 768 on a logarithmic scale, only allowing values divisible by 12; the number of self-attention layers ranged from 2 to 16. The options for the optimizer were stochastic gradient descent, Adam [67], and AdamW [68], and for the scheduler the options were no scheduler, only a warm-up phase, and a warm-up phase followed by a cosine annealing with warm restarts and a linear decay scheduler.

To initiate the hyperparameter optimization process, SMAC3 [66] generated 25 initial random configurations, which were evaluated using the training and validation datasets of the first cross-validation split. The performance of different configurations, measured with the validation loss (mean absolute error (MAE) for supervised models and cross-entropy for masked language model pretraining), guided the selection of subsequent configurations by SMAC3. This iterative process continued until no or negligible improvements in performance were observed. The final hyperparameter configurations for all model types are summarized in Table 1.

**Table 1.** Overview of the hyperparameters selected by SMAC3. The values that were not optimized are shaded in gray. The embedding size and number of layers chosen for the from-scratch model were retained for both pretraining and fine-tuning with the same dataset. During fine-tuning of the pretrained TAPE model, the embedding size and number of layers of the pretrained model were reused.

|  | learning rate | optimizer | scheduler | embedding size | number of layers |
| --- | --- | --- | --- | --- | --- |
| from scratch | 3.12e-5 | Adam | none | 180 | 9 |
| pretraining with own data | 1.94e-4 | Adam | warm-up only | 180 | 9 |
| fine-tuning own | 3.26e-4 | AdamW | warm-up + cosine annealing with decay | 180 | 9 |
| fine-tuning TAPE | 3.11e-4 | AdamW | warm-up + cosine annealing with decay | 768 | 12 |

## Data and code availability

The datasets analyzed during this study and all scripts used to obtain the results are available as open source on GitHub under the Apache 2.0 license at https://github.com/PigeonMark/MTL-peptide-property-prediction and on Zenodo at https://zenodo.org/doi/10.5281/zenodo.11084462. All code is written in Python 3.9 [69]. Pytorch Lightning (version 2.0.1) and PyTorch (version 1.12.0) [70] were used for model development, the model architecture is based on the TAPE (version 0.5) [58] implementation, and SMAC3 (version 2.0.2) [66] was used for hyperparameter tuning. NumPy (version 1.23.5) [71], pandas (version 2.0.0) [72], [73] and scikit-learn (version 1.2.2) [74] were used for data processing and metric calculation. Matplotlib (version 3.7.1) [75] and seaborn (version 0.12.2) [76] were used to create the figures.

## Author contributions

C.D. performed the study. C.D., C.A., and W.B. wrote the manuscript. K.L. and W.B. conceived and supervised the study. K.L. and W.B. revised the manuscript. All authors read and approved the final manuscript.

## Supplementary material

**Supplementary Table 1.** Overview of the ProteomeXchange identifiers of the ProteomeTools project. The data from the ProteomeTools project have been made available through various datasets.

| ProteomeXchange Identifier | Description | Reference |
| --- | --- | --- |
| PXD004732, PXD010595 | Non-modified tryptic peptides | [18] |
| PXD019086 | Non-modified tryptic peptides measured on timsTOF | [11] |
| PXD009449 | Modified tryptic peptides | [19] |
| PXD021013 | HLA class I, HLA class II, and non-tryptic peptides | [23] |
| PXD043844 | HLA class I, HLA class II, and non-tryptic peptides measured on timsTOF | [33] |
| PXD023119 | TMT-labeled non-modified tryptic peptides | [34] |
| PXD023120 | TMT-labeled non-modified non-tryptic peptides | [34] |

**Supplementary Figure S1.**
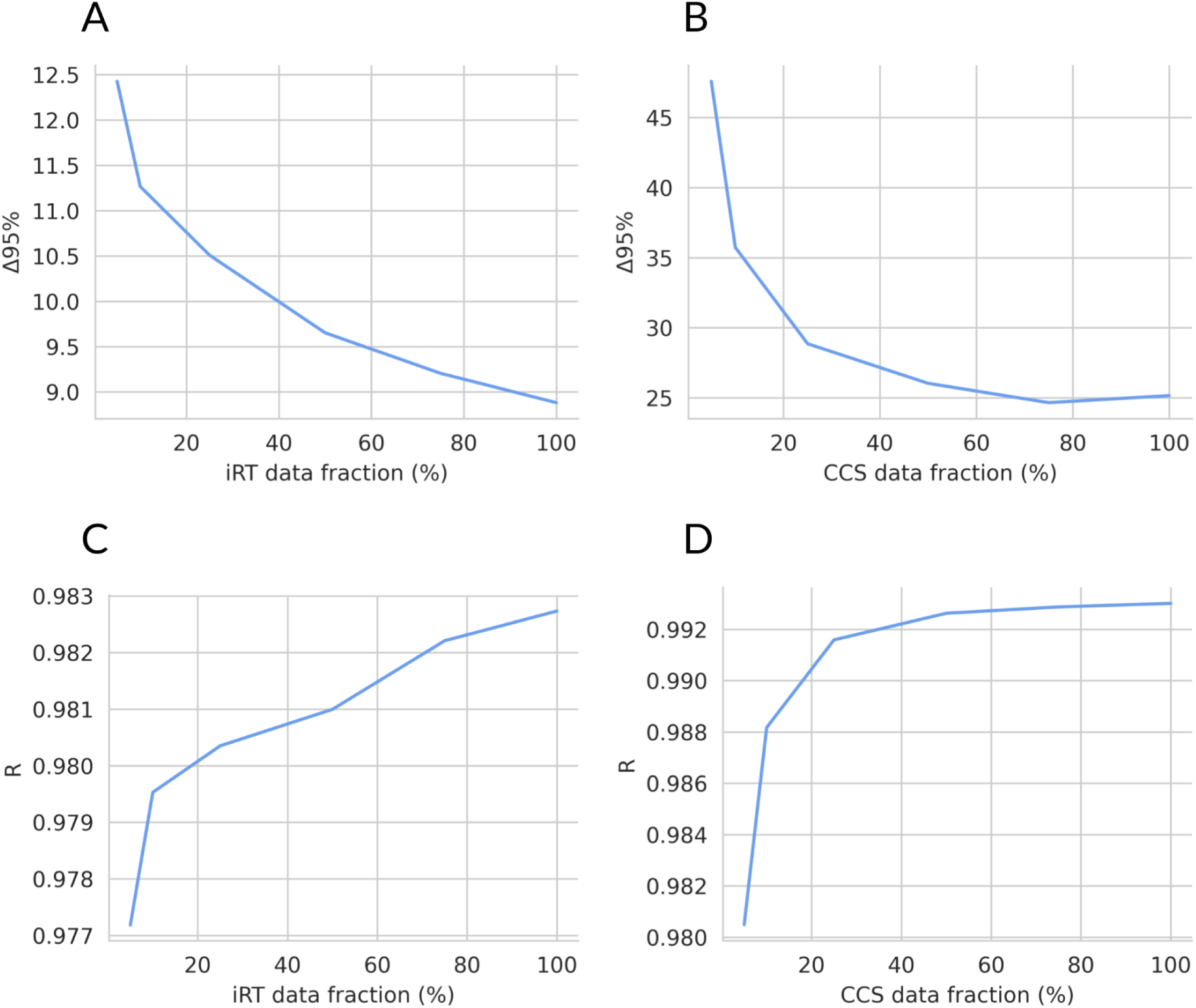
Peptide property prediction learning curves. The performance of a transformer encoder when trained on an increasing fraction of the data measured by the Δ95% of the model predicting the **(A)** iRT and **(B)** CCS of unseen, optionally modified peptides. This metric gives the value so that 95% of the absolute errors are smaller than the Δ95%. Additionally, the performance is measured by the Pearson correlation coëfficient (R) of the model predicting the **(C)** iRT and **(D)** CCS of unseen, optionally modified peptides. 100% of the RT data corresponds to around 1.7 million peptides and 100% of the CCS data is around 539,000 peptides.

**Supplementary Figure S2.**
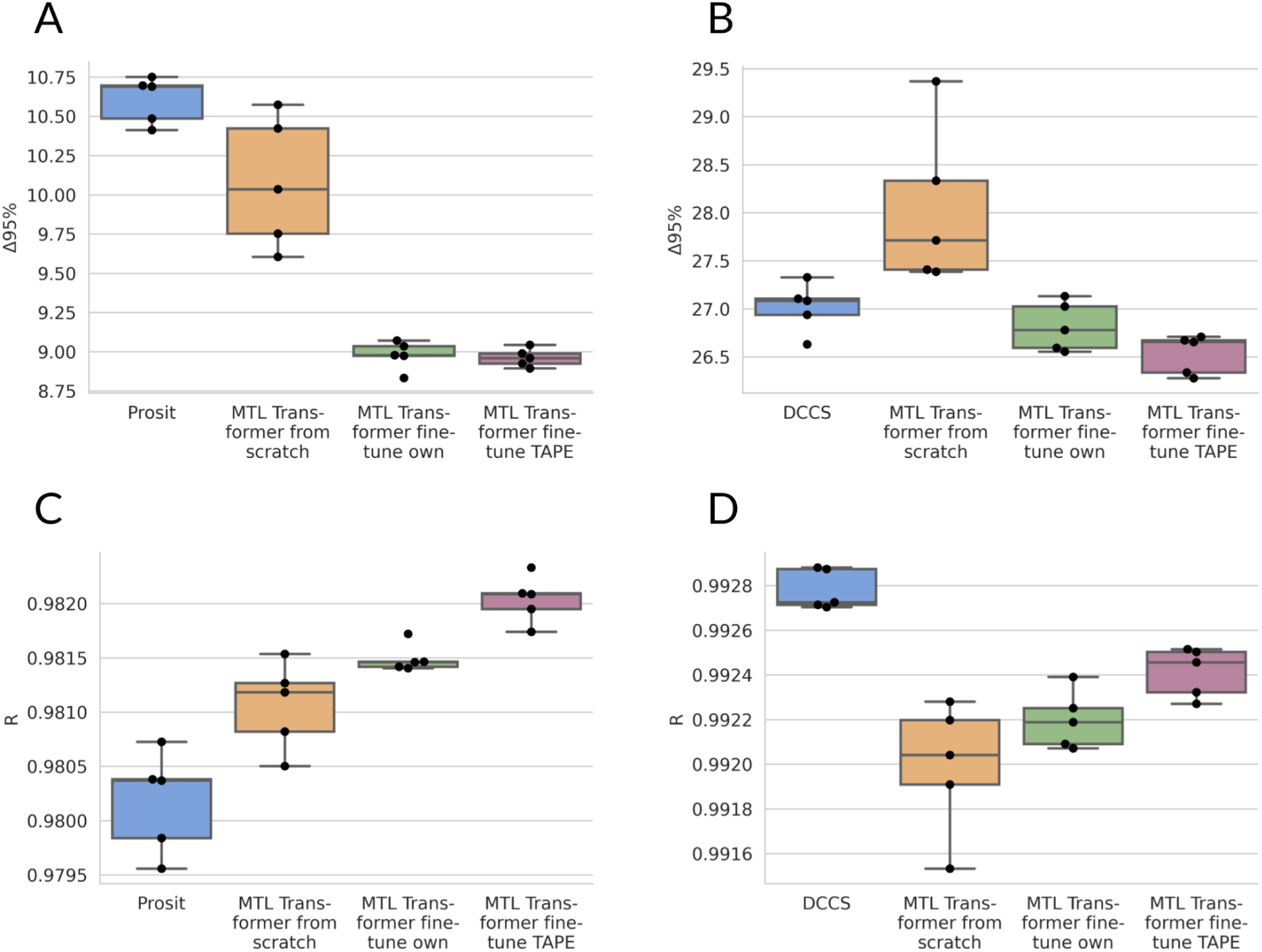
Influence of pretraining on peptide property prediction performance. The five-fold cross-validation performance of multi-task learning (MTL) transformer encoders when trained without pretraining (from scratch), when fine-tuning a model pretrained on the same data (fine-tune own), or when fine-tuning a model pretrained on a large protein dataset (fine-tune TAPE). The performance of a single cross-validation split is measured by the Δ95% of the model predicting the **(A)** iRT and **(B)** CCS of unseen modified peptides. This metric gives the value so that 95% of the absolute errors are smaller than the Δ95%. Additionally, the performance is measured by the Pearson correlation coëfficient (R) of the model predicting the **(C)** iRT and **(D)** CCS of unseen, optionally modified peptides. The performance of Prosit [4] for iRT prediction and DCCS [11] for CCS prediction, retrained and evaluated on the same cross-validation splits, is also given as reference. Boxplots are constructed as follows: the box extends from the lower to upper quartile values of the data, with a line at the median. The whiskers extend from the box to the last datum before 1.5 times the interquartile range above/below the box.

**Supplementary Figure S3.**
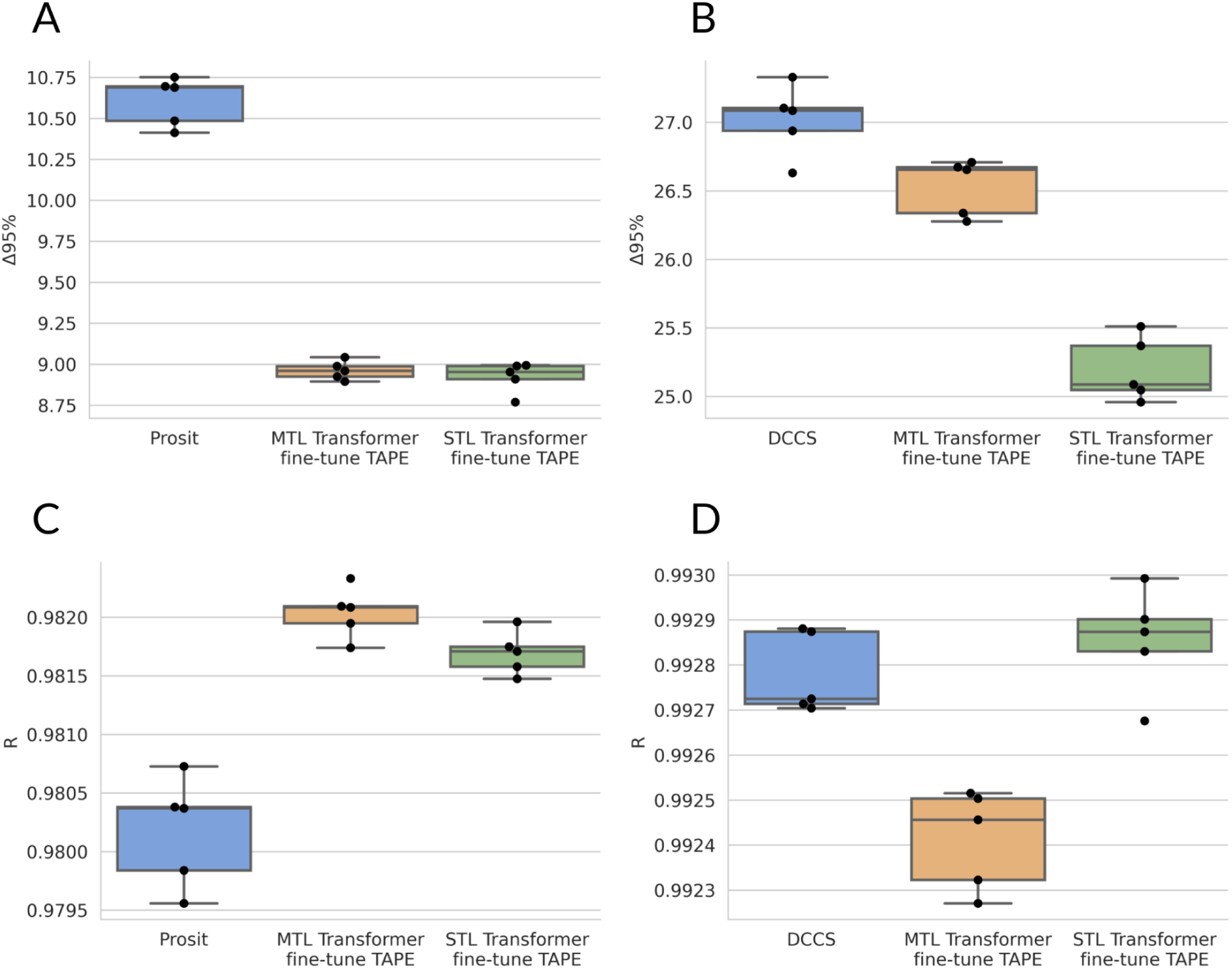
Influence of multi-task learning on peptide property prediction performance. The five-fold cross-validation performance of a multi-task learning (MTL) and a single-task learning (STL) transformer encoder. The performance of a single cross-validation split is measured by the Δ95% of the model predicting the **(A)** iRT and **(B)** CCS of unseen modified peptides. This metric gives the value so that 95% of the absolute errors are smaller than the Δ95%. Additionally, the performance is measured by the Pearson correlation coëfficient (R) of the model predicting the **(C)** iRT and **(D)** CCS of unseen, optionally modified peptides. The performance of Prosit [4] for iRT prediction and DCCS [11] for CCS prediction, retrained and evaluated on the same cross-validation splits, is also given as reference. Boxplots are constructed as follows: the box extends from the lower to upper quartile values of the data, with a line at the median. The whiskers extend from the box to the last datum before 1.5 times the interquartile range above/below the box.

## Footnotes

1 https://massive.ucsd.edu/ProteoSAFe/static/massive-kb-libraries.jsp

2 https://github.com/searlelab/chronologer/tree/main/data

